# Genome-wide selection footprints and deleterious variations in young Asian allotetraploid rapeseed

**DOI:** 10.1101/412551

**Authors:** Jun Zou, Lingfeng Mao, Jie Qiu, Meng Wang, Zhesi He, Lei Jia, Dongya Wu, Yongji Huang, Meihong Chen, Yifei Shen, Enhui Shen, Ruiyuan Li, Dandan Hu, Kai Wang, Lei Shi, Chuyu Ye, Ian Bancroft, Graham J King, Jinling Meng, Longjiang Fan

**Affiliations:** National Key Laboratory of Crop Genetic Improvement, Huazhong Agricultural University, Wuhan 430070, China; Institute of Crop Sciences & Institute of Bioinformatics, Zhejiang University, Hangzhou 310058, China; Department of Biology, York University, Heslington, United Kingdom; Center for Genomics and Biotechnology, Haixia Institute of Science and Technology (HIST), Fujian Agriculture and Forestry University, Fuzhou, Fujian 350002, China; Southern Cross Plant Science, Southern Cross University, PO Box 157, Lismore NSW 2480, Australia

**Keywords:** Allopolyploid, selection footprints, deleterious variations, introgression, rapeseed

## Abstract

*Brassica napus* (AACC, 2n=38), is an important oilseed crop grown worldwide. However, little is known about the population evolution of this species, the genomic difference between its major genetic clusters, such as European and Asian rapeseed, and impacts of historical large-sale introgression events in this young tetraploid. In this study, we reported the *de novo* assembly of the genome sequences of an Asian rapeseed (*B. napus*), Ningyou 7 and its four progenitors and carried out *de novo* assembly-based comparison, pedigree and population analysis with other available genomic data from diverse European and Asian cultivars. Our results showed that Asian rapeseed originally derived from European rapeseed, but it had subsequently significantly diverged, with rapid genome differentiation after intensive local breeding selection. The first historical introgression of *B. rapa* dramatically broadened the allelic pool of Asian *B. napus*, but decreased their deleterious variations. The secondary historical introgression of European rapeseed (canola-quality) has reshaped Asian rapeseed into two groups, accompanied by an increase in genetic load. This study demonstrates distinctive genomic footprints by recent intra- and inter-species introgression events for local adaptation, and provide novel insights for understanding the rapid genome evolution of a young allopolyploid crop.

## Introduction

Advances in crop performance attributable to genetic improvement during the ‘Green Revolution’ were achieved using a range of plant breeding approaches that often differed between crops and major geographical regions. Although yield increases were in general attained during a period of atypical climate stability (Pingali, 2012), there is now increasing variability with significant local variation in microclimate as well as seasonal meteorological patterns that affect crop phenology. In order to develop climate-resilient crops for the coming decades there is a need to understand the consequences of different historical breeding strategies (Kole *et al*., 2015). In particular, there is opportunity to establish the value of allelic pools that traditionally have contributed to locally adapted landraces following initial domestication and subsequent radiations of crop species. Polyploidy has played an important role throughout plant evolution, and due to locus redundancy and processes such as neo-functionalisation and sub-functionalisation of paralogous genes, can provide a platform for developing resilient crops (Roulin *et al*., 2013). *Brassica* rapeseed presents a unique diagnostic system for studying such processes.

*Brassica napus* L. (A^n^A^n^C^n^C^n^, 2n=38) is globally the third largest source of vegetable oil and plays an important role in wheat-based arable systems (Ebrahimi *et al*., 2017). This tetraploid rapeseed is a very young allopolyploid crop derived from interspecific crosses between the diploid progenitors, *B. rapa* (A^r^A^r^, 2n=20) and *B. oleracea* (C°C°, 2n=18). Although having a short history of post-Neolithic speciation (~7,500 years) (Chalhoub *et al*., 2014) and domestication (~700 years) (Chalhoub *et al*., 2014), it interestingly displays relatively rapid LD decay rates, especially in the A genome where it is less than 80 Kb (Qian *et al*., 2014). This compares with other allopolyploid crops such as cotton and wheat that have also had a domestication history of over 4,000 years (Peng *et al*., 2011), but which display relatively slow LD decay rates (at approximately 700 Kb (Ma *et al*., 2018; Wang *et al*., 2017a) and ~4cM (Gurung *et al*., 2014), respectively).

No wild populations of *B. napus* have been found to date, making it challenging to determine the species’ origins. Nonetheless, deducing the origin and unravelling the impact of secondary radiation in recent domestication on genome organization and allelic selection for *B. napus* is of intrinsic and practical value (Becker *et al*., 1995; Wang *et al*., 2014).

Several distinct genetic groups of *B. napus* oilseed cultivars have been classified according to their geographic origin and seasonal crop type, including European winter, Asian semi-winter, Canadian, Australian and European spring types. In China, the local *B. rapa* oilseed crop had been cultivated for more than a thousand years (Li, 2001), but from the 1950s this crop was extensively replaced by *B. napus* “Shengliyoucai” (SL) types, which included a series of cultivars and derived lines that had been developed following selective breeding (Liu, 2000). Many distinct cultivars were further derived based on SL with introgression of *B. rapa*, an approach frequently adopted to broaden the genetic diversity and improve the local adaptation of *B. napus* in Asia (Zhang *et al*., 2014). As a result, many Asian cultivars carry a common lineage derived from SL and the *B. rapa* introgression. In the 1970s, Asian rapeseed was further significantly improved by the introgression of European ‘double low’ alleles which had been discovered in Canada and then re-introduced to European rapeseed lines, providing low seed glucosinolate and erucic acid.

Dissecting the genome of an Asian rapeseed cultivar with SL pedigree and *B. rapa* introgression in the absence of “double low” introgression provides the opportunity to understand the evolutionary footprints associated with local adaptation and artificial selection, and identify footprints of genome differentiation compared with modern European *B. napus*. In this study, we *de novo* sequenced and assembled Ningyou 7 (NY7), which is an elite semi-winter cultivar carrying mid-20^th^ century introgressions of local diploid rapeseed *B. rapa* from the early SL cultivars in China prior to extensive introgression of “double low” traits from European cultivars (Luo *et al*., 2017; Qiu *et al*., 2006) (Figure S1, more details see Methods). This contrasts with the recently sequenced Asian cultivar ZS11 which carries double low alleles (Sun *et al*., 2017). A comprehensive suite of comparisons was undertaken to unravel contributions to the genome of NY7 from its four ancestors, and contrast with the available genome sequences of diverse European and more modern Asian rapeseed cultivars (Huang *et al*., 2013; Schmutzer *et al*., 2015; Shen *et al*., 2017; Wang *et al*., 2018). The *de novo* assembled pedigree and comparison to *B. napus* genomes from other lineages helped us trace the origins of valuable adapted trait alleles, and revealed which genomic structural variations had been selected during recent Asian rapeseed breeding. These results provided novel insights into the rapid evolutional signatures and significant impacts of intra-specific and inter-specific introgression, arising from local selective breeding of a young allopolyploid species.

## Results

### *De novo* genome assemblies of NY7

We sequenced the genome of the Asian *B. napus* cultivar NY7 (pedigree see Figure S1) combining three technologies (Table 1; details see Table S1). The assembled genome based on Illumina and Pacbio data was further confirmed by making use of the syntenic relationship of an updated *Bna*TNDH genetic linkage map constructed using 353 DH lines with a total of 2,304 backbone markers (Table S2). The resulting assembly spans 994 Mb, covering ~85% of the genome size based on *K*-mer estimation of the NY7 genome size (1170 Mb, Figure S2). The assembly was further improved by three-dimensional (3D) chromosome conformation capture sequencing (HiC), increasing the scaffold N50 to 6.90 Mb (Table 1, Figure S3). Based on this assembly, we generated 19 chromosomal pseudomolecules anchored to the updated *Bna*TNDH genetic linkage map and two other published genetic linkage maps as reference (Chalhoub *et al*., 2014; Delourme *et al*., 2013) (Figure S4). 372 scaffolds were anchored to the linkage maps, representing 890 Mb (89.5%) of the total assembly length (Figure 1a; Table 1, Figure S5). This represents a more comprehensive coverage of the genome compared with Darmor-*bzh* and ZS11 (Table 1).

**Table 1.** | Summary of genome assembly and annotation of Asian rapeseed NY7 and its pedigree

| **NY7** |  |  |  |  |
| --- | --- | --- | --- | --- |
| Sequencing | Insert libraries | Illumina | Pacbio RS II | HiC |
|  | 160 bp-20 Kb | ~200× | ~30× | ~100× |
| Scaffolds | N50 size (Mb) | L50 | The maximum (Mb) | Total non-N size (Mb) |
|  | 1.27 | 229 | 7.41 | 956.9 |
| Super-scaffolds by HiC | 6.91 | 37 | 33.78 | 956.9 |
| Chromosomes by two genetic maps | Total anchored /non-N size (Mb) | Unique marker No. | **Darmor-*bzh*<sup>a</sup>**<br>Anchored/non-N size | **ZS11<sup>b</sup>**<br>Anchored/non-N size |
|  | 892.0/874.0 | 13,164 | 645.4/553.4 | 855.0/797.7 |
| Annotation | Gene models | BUCSOs/CEGMA (%) | Supported by EST and Swiss-prot etc. (%) | Repetitive elements (%) |
|  | 104,179 | 98.5 / 98.4 | 97.0 | 45.0 |
| **Parental lines in NY7 pedigree** |  |  |  |  |
| Parental lines* | 160 bp-10 Kb Illumina data (Gb)<sup>#</sup> | *K*-mer estimate size (Gb) | Genome assembly (Mb) | Scaffold N50 size (Kb) |
| SL-1 | 147.4 | 1.175 | 860.7 | 331.0 |
| CDA | 106.6 | 0.514 | 412.2 | 178.9 |
| NY1 | 153.0 | 1.221 | 845.7 | 369.6 |
| CY2 | 118.1 | 1.178 | 869.5 | 293.2 |

\*details see Figure 1a
<sup>#</sup>including the libraries with 3 and 10 Kb insertion sizes.
<sup>a,b</sup> Comparison of assemblies indicates that whilst NY7 included only 2% (18Mb) as ‘N’, Darmor-*bzh* included 14% (92Mb) and ZS11 7% (57Mb).

**Figure 1.**
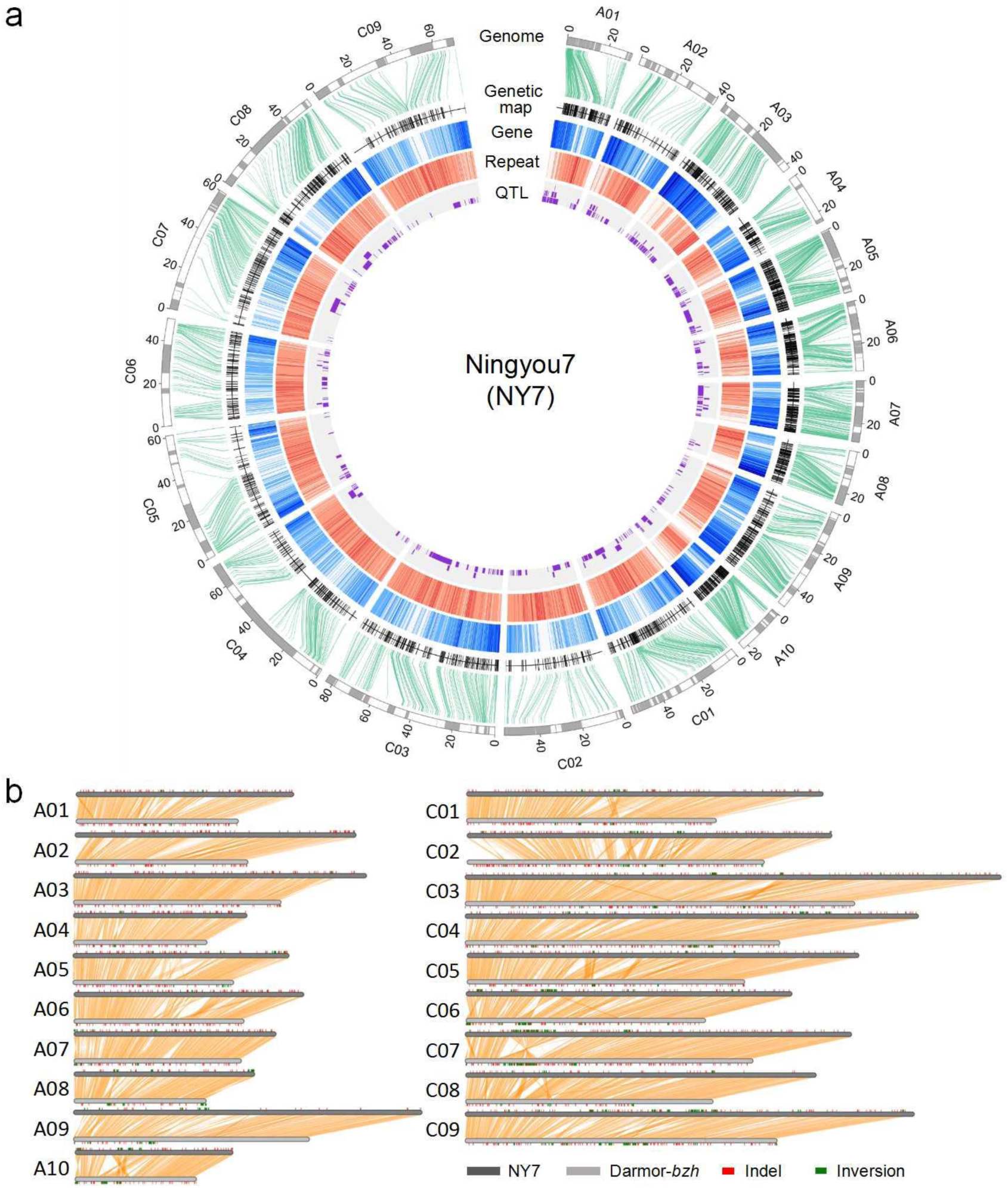
The genome assembly of NY7. (a) Circos plot of the NY7 genome assembly with genetic map, annotation and QTLs. Circles inwards: chromosome with linked scaffolds (a scaffold with a same color and linked scaffolds with different corlors), genetic maps, gene density, repeat density and QTLs based on NY7 and Tapidor (*Bna*TNDH) population. (b) Genomic synteny between two *de novo* assembly genomes, NY7 and Darmor-*bzh*.

To evaluate the quality of our NY7 genome assembly, 643,944 publicly available *B. napus* expressed sequence tags (ESTs) from GenBank were mapped to the genome, of which ~97% could be aligned with an average identity of 97%. We further examined the assembly based on the congruence of transcriptome genotypes of 47 *Bna*TNDH lines and built genome-ordered graphical genotypes (GOCGs) (Table S3). This allowed us to include an additional ~1.3 Mb (two scaffolds) as well as correct 52 mis-assemblies with cumulative length of ~15.9 Mb into the 19 pseudo-chromosomes (details see Methods; Table S3).

As found in other *B. napus* genome assemblies, the NY7 genome includes a number of extensive repetitive regions in which no or few molecular markers could be found (Figure 1a). For example, by using HiC sequence data (Figure S6) and centromeric-specific markers from the *Bna*TNDH genetic map (Long *et al*., 2011), we were able to extend the end of A03 in the NY7 assembly by about 3.7 Mb (38.3~42.0 Mb), whereas it had previously been unresolved by anchoring in the three previously sequenced *B. napus* genomes (Bayer *et al*., 2017; Chalhoub *et al*., 2014; Sun *et al*., 2017). These genomic regions were not found within the anchored Darmor-*bzh* pseudo-chromosomes but in an unanchored scaffold that had been assigned to chromosome A03 of Darmor-*bzh* and other *Brassica* A03 genomes such as *B. rapa* cv. Chiifu-401 v2.5 (Wang *et al*., 2011).

A total of 104,179 protein-coding genes were annotated (Table 1). The assembly appears to cover the majority of the gene space (Table S4) based on evidence of 98.5% of CEG dataset genes mapping to NY7 genes and 98.4% of the plantae BUSCO dataset (V3) (Simao *et al*., 2015). Repetitive sequences accounted for 44.7% of the genome assembly (Table 1), with 16.2% of the genome represented by long terminal retrotransposons.

### Structural variation between Asian and European cultivars based on *de novo* assembly

We carried out pairwise comparison of NY7 with the sequenced genomes of European cultivars Darmor-*bzh* and Tapidor, and Asian cultivar ZS11, all of which carry “double low” introgressed alleles at multiple loci, focusing on the comparison between NY7 and Darmor-*bzh*. Though there was a high level of synteny, frequent small and large structural variations, along with common structural variations were observed (Figure 1b; Figure S6, Table S5-6). This included many small and large present and absent variations (PAV) between NY7 and Darmor-*bzh.* With NY7 as reference, 1.49 million SNP and 0.17 million small insertion/deletions (Indels) were observed between the two cultivars. When the Darmor-*bzh* genome was compared to the NY7 reference genome, there were rich large structural variations, such as 34.5k of inversion events covering 45.1Mb (26 bp~100 Kb in size), 4.4k translocation events covering 9.4Mb (>1 Kb in size), 42.9k insertions covering 118.3Mb, and 82.3k deletions covering 162.1Mb (Table S5, Figure 1b). An increased frequency of insertion events could be explained by the larger assembled genome size of NY7. 10.0 Mb of PAV segments were identified in NY7, similar to 10.9 Mb in Darmor-*bzh*, including 110 large PAV > 5 Kb in NY7, and 91 in Darmor-*bzh* (Table S6).

In order to compare NY7 to the other parent of the *Bna*TNDH population, we generated an improved *de novo* genome assembly for the double-low European cultivar Tapidor, which had previously been *de novo* assembled (Bayer *et al*., 2017) based on large-insert library sequencing data (Table S1; the genome assembly of Tapidor with an accumulated 1001.7 Mb in length and a 826.6 Kb scaffold N50 size). We then anchored QTL to the two parental genomes that accounted for seed yield and its related traits identified in the *Bna*TNDH population with prediction accuracy above 0.5. At least 25 QTL were located in regions with PAV (Table S7). For example, within a 2.2 Mb QTL region (*es. C3-3*) (Figure S7), Tapidor had an average coverage depth of only 3.0 compared to the average depth of 15, and just 3.6% of orthologous sequence could be aligned to the corresponding QTL region in NY7.

### Phylogeny and genetic diversity of Asian and European populations

To analyze the genetic differentiation of Asian *B. napus* cultivars from European cultivars, genomic data from 68 Asian (30 double-low and 38 double-high) (Chalhoub *et al*., 2014; Huang *et al*., 2013; Shen *et al*., 2017; Wang *et al*., 2018) and 59 European representative accessions (Chalhoub *et al*., 2014; Schmutzer *et al*., 2015; Wang *et al*., 2018) generated in this and previous studies were collated (Table S8; including three additional SL derived early cultivars. See Methods). Phylogenetic and principle component analysis (PCA) (Figure 2, Figure S8) showed two distinct independent branches representing the Europe and Asian lineages. While the ‘double-low’ subgroup of Asian rapeseeds was clearly separated from the ‘double-high’ subgroup, the former was genetically closer to the European lines (Figure 2). These results suggest that the Asian rapeseed genomes have been re-shaped by strong local selection in a relative short time. This may have resulted in emergence of a distinct domesticated ecotype with a unique and identifiable genetic background.

**Figure 2.**
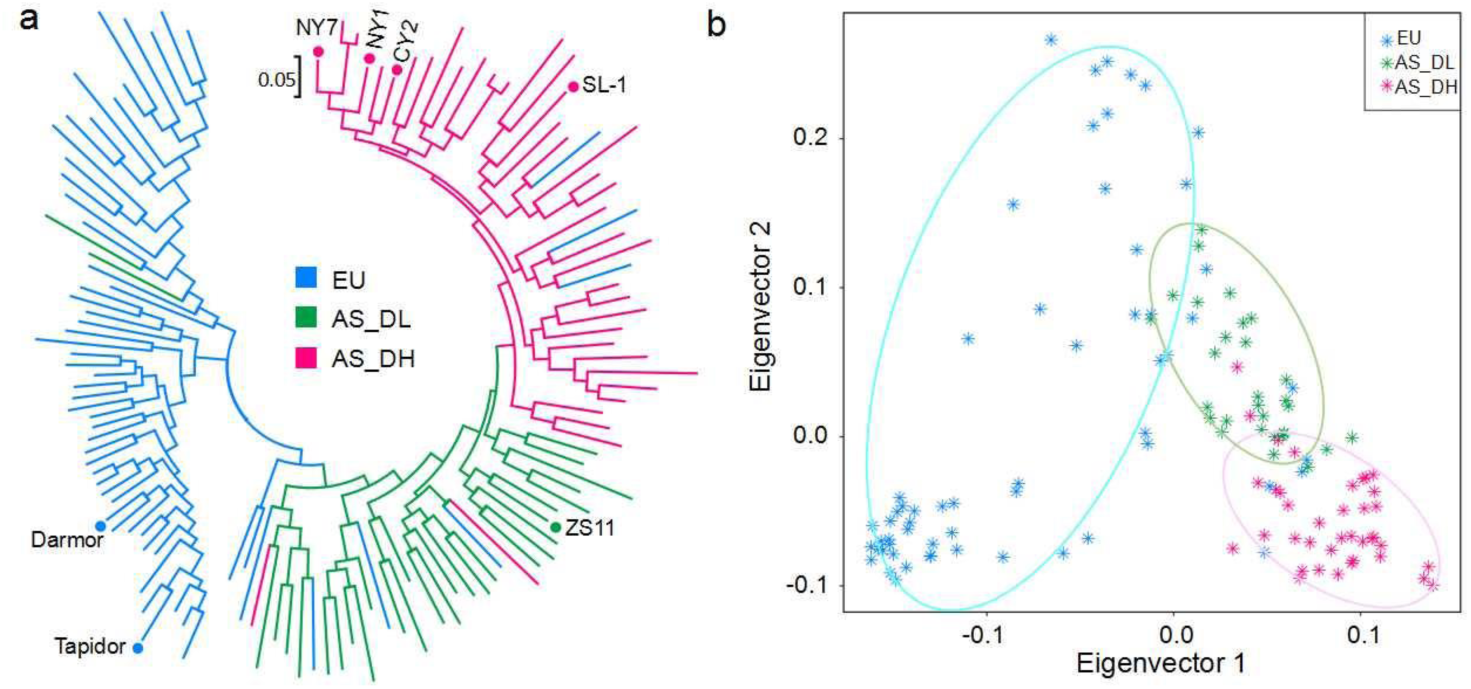
Genetic divergence of Asian and European rapeseeds. (a) Phylogenetic tree and (b) Principle component analysis of Asian (AS) and European (EU) rapeseeds. Asian rapeseeds include double-high (AS_DH) and double-low (AS_DL) varieties. The sequenced cultivars show with stars.

We calculated whole-genome genetic diversity (π) for each subgroup, and found that the European group encompassed the highest diversity (1.45e-3), followed by the subgroups of Asian double-low (1.41e-3) and double-high (1.19e-3). The extensive diversity in the European group is particularly apparent in the C subgenome (Figure S9). For the A subgenome, diversity was highest within the double-low group (1.89e-3), followed by the double-high (1.79e-3) and European groups (1.78e-3). Our analysis of linkage disequilibrium (LD) indicated that double-low Asian rapeseeds have the slowest LD decay rate in both AA and CC subgenomes, while the most rapid decay rate was observed amongst the European cultivars (Figure S10).

### Deducing demographic origins of Asian rapeseeds

To determine the origin of Asian rapeseeds, demographic analysis was carried out based on the available SNP data, including data from germplasm representing the two domesticated diploid progenitor species (34 *B. rapa* and 37 *B. oleracea*) (Cheng *et al*., 2016). We attempted to unravel the origin of Asian rapeseeds using four different demographic simulation models (Figure S11) based on putative neutral SNPs in intergenic regions (details see Methods). Demographic analysis based on the AA subgenome supported the model that the European *B. napus* AA subgenome (*Bna*A) first diverged from its ancestor *B. rapa* and subsequently bifurcated to generate Asian rapeseed populations (Figure 3a). A similar outcome was observed based on the CC subgenome (Table S9). In combination, these results support a European origin for Asian rapeseeds, i.e. Asian rapeseeds were introduced from Europe, which is consistent with historic records (Liu, 2000). The timing of divergence between the European and Asian groups was further estimated using SMC++ (Terhorst *et al*., 2017). Based on the 9e-9 substitute rate (Qi *et al*., 2017), Asian rapeseeds appear to have split from the European group within the past 500 years (Figure 3b), with a notable decrease in population size representing a genetic bottleneck. In order to investigate the origin of Asian double-low rapeseeds further, the same demographic analysis approach was performed to test different origin models (Figure S12). We found evidence that the hybridization origin model (Figure 3a. i.e. double-low first split from double-high Asian, with introgression from European double low rapeseeds) best fits the genomic data (Table S9), consistent with historical records (Liu, 2000).

**Figure 3.**
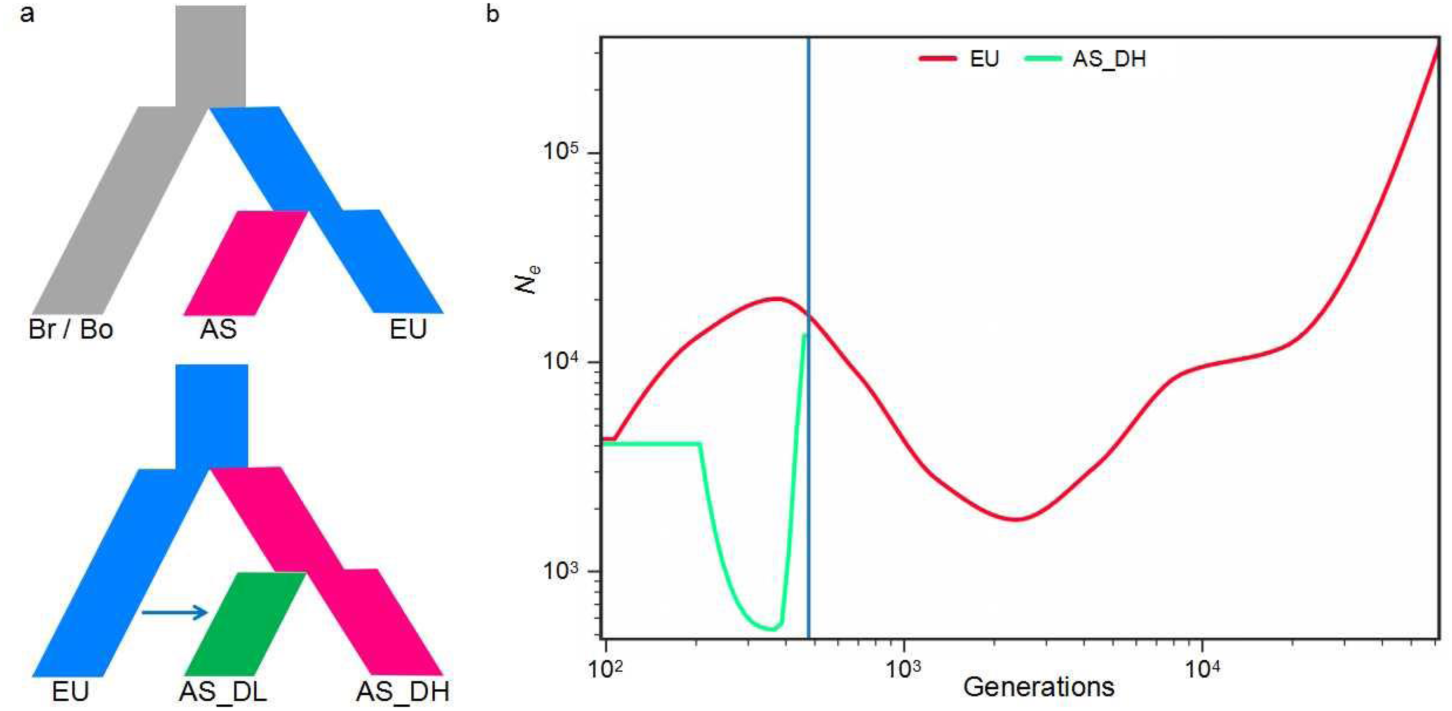
Demography inference for Asian rapeseeds. (a) The best demographic models for the origin of Asian and its double-high (AS_DH) and double-low (AS_DL) varieties. EU, Bo, and Ba refer to European rapeseed and its two progenitors, *B. rapa* and *B. oleracea*. (b) Divergent time (generation) and effective population size (Ne) changes for European and Asian rapeseed populations using SMC++.

### Genomic signatures of Asian rapeseeds under local selection

We performed large-scale genomic scans (*F*_ST_) between double-high/-low Asian and European rapeseeds, in order to uncover the most differentiated genomic regions between European and Asian rapeseed populations, and to reveal genomic signatures of the breeding processes associated with Asian rapeseeds. Between the double-high Asian and European groups, a total of 1,665 highly differentiated scanning windows (Z(*F*_ST_) > 3) were found, covering 28.3 Mb of the genome and containing 2,519 genes (Figure 3). An asymmetric subgenome selection pattern was found, with a larger size in the CC (23.1 Mb) than the AA (5.2 Mb) subgenome. Previously characterized QTLs for important traits detected in the *Bna*TNDH population reside within these highly divergent regions (Table S10). For example, a candidate selected genomic region detected in A09 (40.0~40.8 Mb) overlapped with a QTL (*es. A9-32*) associated with maturity time and development time of seed, co-locating with an orthologue of a flowering time controlling gene *agl17-2* (Figure 4a). In addition, other flowering time related QTLs (*es. C7-16*, *es. C7-17*) could also be found within the genomic regions highly divergent between the European and Asian groups.

**Figure 4.**
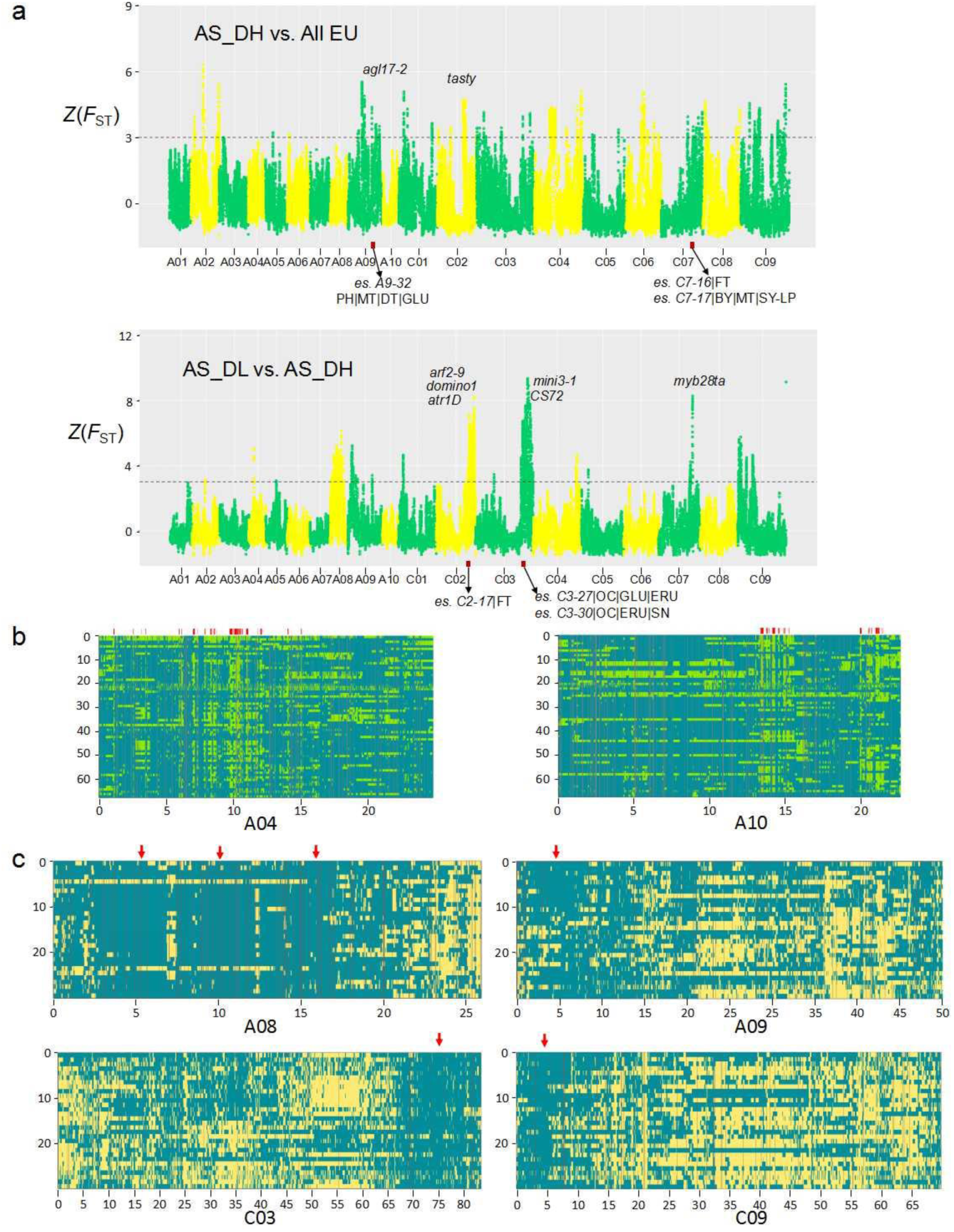
Genomic selection signals in Asian rapeseeds. (a) Significantly divergent genomic regions of Asian double-high rapeseeds (AS_DH) relative to the European rapeseeds (EU) and Asian double-low rapeseeds (AS_DL) relative to AS_DH. Some genes and QTLs locating in those regions were indicated. (b) Local ancestry inference of the introgression of *B. rapa* for Asian rapeseed shown with A04 and A10. On the top of the box, the red signals present the conserved blocks with introgression of *B. rapa* (green). (c) Local ancestry inference of the introgression of European lineage (blue) in Asian double low cultivars (AS_DL). T genomic regions (with arrow) of AS_DL in four chromosomes illustrated still keep EU genetic background up to now even undergone strong local selection.

Highly divergent genomic selection signatures were also detected between the double-low and double-high rapeseeds, covering 30.2 Mb genomic regions (AA: 8.0 Mb; CC: 22.2 Mb). Five Mb-scale peaks found in A08, A09, C02, C03 and C09 perfectly matched results from a previous GWAS study (Wang *et al*., 2018), where genes were identified associated with seed-quality traits, including erucic acid content (EAC), glucosinolate content (GSC) and seed oil content (SOC). Two QTLs (*es. C3-27* and *es. C3-30*) which contribute to these traits also locate in the highly divergent region in chromosome C03 of the *Bna*TNDH population (Figure 4a, Table S7). These genomic footprints provide insights into the recent genomic footprints of ‘double-low’ rapeseed breeding for Asian rapeseeds.

We further performed a local ancestry inference (LAI) analysis to trace the potential introgressed regions of European *B. napus* and diploid *B. rapa* within Asian rapeseeds. Interestingly, there were extensive introgression signatures of *B. rapa* in Asian rapeseeds, with a highly conserved region covering 8.8 Mb containing important allelic patterns maintained in the Asian rapeseed population. This includes alleles for *BnFLC.A10* that are distinct from those found in European cultivars (Figure 4b). There is strong evidence that alleles from *B. rapa* that contribute to early flowering time, high erucic acid, *Sclerotinia* resistance, and yield traits, became fixed in the gene pool of Asian double high rapeseeds (Table S11). In contrast, the highly conserved regions introgressed from European cultivars into Asian “double-low” cultivars, notably those on A08, A09 and C03, contain well characterized alleles for major genes accounting for low glucosinolate and erucic acid that are prevalent in the modern rapeseed gene pool (Figure 4c).

### Genomic footprints of breeding on the NY7 pedigree and Asian cultivars

In order to understand the pattern of inheritance contributing to NY7, we also sequenced four parental lines (SL-1, CDA, NY1, CY2) from the NY7 pedigree, including the founder landrace SL and the *B. rapa* CDA (Table 1 and Figure S1). Drafts of the four genomes were *de novo* assembled by deep sequencing with libraries of different 3-10 Kb insertion sizes (Table S1) with final genome assemblies (an average 300 Kb of scaffold N50 size) covering 70~80% of estimated genome sizes (Table 1).

Identity by descent (IBD) of the NY7 pedigree was estimated to determine patterns of inheritance (Figure 5). As expected, the greatest overall genetic contributions to NY7 were attributable to the two most recent parental lines NY1 (40%) and CY2 (46%) (Figure 5), consistent with previous results estimated with 29,347 SNP markers covering 643.7Mb (Wang *et al*., 2017b). Meanwhile, a greater proportion of genomic segments originating from progenitor parental lines SL (at least 8.41%) and CDA (at least 5.8%) remained and were detectable in the AACC genomes of cultivar NY7 than estimated by the earlier SNP study, evidence of retention following successive rounds of selection in the breeding program (Figure 5; a detailed table of IBD regions with QTL information see Table S12).

**Figure 5.**
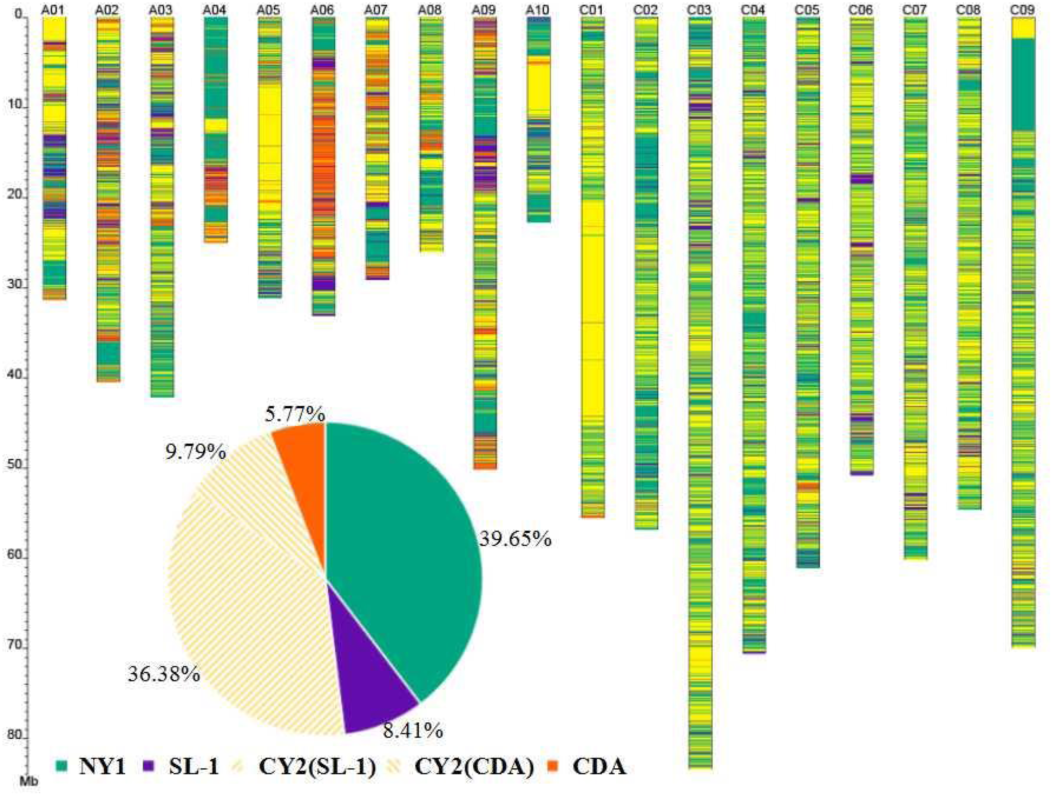
Identity by descent (IBD) inheritance pattern for the NY7 pedigree. The genomic contributions by four different parental lines to NY7 are coded in chromosomes as CY2 (yellow), CDA (orange), NY1 (green), and SL1 (purple). The genomic contributions of SL1 and CDA to CY2 when using CY2 as reference was also divided with the same yellow color as CY2. Accumulated percentages of contribution by the four parental lines were shown in a circle in the figure.

We were able to resolve many contributions from large-scale contiguous Mb-scale genomic blocks that had been inherited intact from one of the parental lines into NY7, and which could not be estimated by SNP markers in previous studies. For example, a ~25 Mb region (position of 20 ~ 45 Mb) in the middle of chromosome C01 of NY7 was mostly contributed by CY2 (Figure 5). Meanwhile, almost no assembled scaffolds or reads of NY1 could be mapped within this region in the NY7 reference, indicating that the 25 Mb-scale region originally from the C01 chromosome was completely absent in NY1. This may have been caused by homoeologous exchange between A01 and C01, which has been frequently observed in other studies (Chalhoub *et al*., 2014; Xiong *et al*., 2011). A further example from 0~12.6 Mb at the top of C09 of NY7 indicated that the segment 0~2.3 Mb was completely absent in NY1, with the remainder absent in CY2.

SL is a winter cultivar used as a female parent in a hybridization with an Asian rapeseed *B. rapa* line (here CDA) that led to establishment of a semi-winter cultivar adapted to Asian production areas. CDA contributed at least 5.8% genomic regions to NY7 based on IBD estimation (Figure 5), where favorable alleles of QTLs that contribute to maturation time (*es. A1-24*), oil content (*es. A4-19*, *es. A6-10*), seed weight (*es. A4-4*), and disease resistance (*es. A4-15*) are contributed by CDA (Table S12). Within this region, we found a number of orthologues to genes with functions involved in early flowering such as *SEF*, *ELF8*, *REF6*, and *VIP4* (Table S12). It is worth mentioning that the contribution of the CDA genome (5.8%) was initially underestimated, as CDA is also one of the donors for NY7’s direct parent CY2. Therefore, a further contribution of CDA was subtracted from CY2, contributing a further 9.79% (Figure 5).

We further examined whether any alleles derived from Chinese *B. rapa* may have remained under selection in the Asian rapeseed population. We focused on non-synonymous SNPs in the Chinese *B. rapa* accession CDA compared to European accessions, and investigated the allele frequency of those SNPs within the flowering related genes of Asian and European populations. We found the gene with the most differentiated allele frequency between the double-high and European groups is a *HDA5* homolog (chrA02g004876), in which the allele frequency is 0.14 in European lines, but above 0.80 in all Asian groups. Notably, four other flowering-time related genes (*PRR5*, *HUA2*, *VIP4*, *CDF1*) reside nearby (A02: 37.8~39.1 Mb) each with an increased allele frequency and almost fixed in Asian groups (Table S13). In addition to the marked changes in allele frequency, we found a present and absent variation (PAV) for a *CDF2* homologue, which could only be completely mapped by sequence reads from CY2 and NY7 but not by reads from SL, and NY1 (Figure S13). The *CDF* (*CYCLING DOF FACTOR*) gene encodes a repressor of the *CONSTANS* (*CO*) transcription factor, which is a key regulator of the photoperiodic flowering response in Arabidopsis (Imaizumi *et al*., 2005). It supported that local introgression of *B. Rapa* contributed many beneficial alleles or genes for NY7 and other Asian cultivars by conferring a shift to semi-winter phenotype under local selection in China.

### Variation of genetic load in the pedigree of NY7 and Asian rapeseeds

Several studies have suggested the presence of a “cost of domestication”, because crops may harbor deleterious mutations that reduce their relative fitness (Eyre-Walker and Keightley, 2007; Loewe *et al*., 2006). To understand better the dynamic changes of deleterious SNPs (dSNP) frequency in Asian rapeseed populations during the different phases of breeding improvement, the dSNP allele frequency for three rapeseed populations (EU, AS_DH, AS_DH) was estimated. In general we found distinct frequency patterns attributable to the AA and CC subgenomes. In the CC subgenome, the AS_DL group has accumulated more fixed dSNPs than the AS_DH or EU group (Figure S14). Interestingly, however, a much lower percentage of fixed dSNPs was observed in the double-high Asian group within the AA subgenome (Figure 6). Furthermore, we found that the relative frequency of deleterious to neutral variants (intergenic SNPs) is clearly lower in the Asian double-high rapeseed population than in the progenitor European population for the AA subgenome. However, this frequency subsequently increased again in the Asian double-low rapeseeds (Figure 6). To examine the effect of *B. rapa* introgression to dSNP accumulation directly, we further estimated the percentage of dSNPs across the individual genomes of NY7 pedigree members. Within the pedigree, CY2 had a sudden reduction in dSNP percentage compared to its immediate predecessor SL in both AA and CC subgenomes. The dSNP percentage progressively increased again for NY7, possibly due to domestication cost associated with breeding programs (Kono *et al*., 2016; Liu *et al*., 2017; Marsden *et al*., 2016). We infer that the *B. rapa* hybrid event or introgression has greatly contributed to slowing the subsequent accumulation of dSNPs in Asian rapeseed.

**Figure 6.**
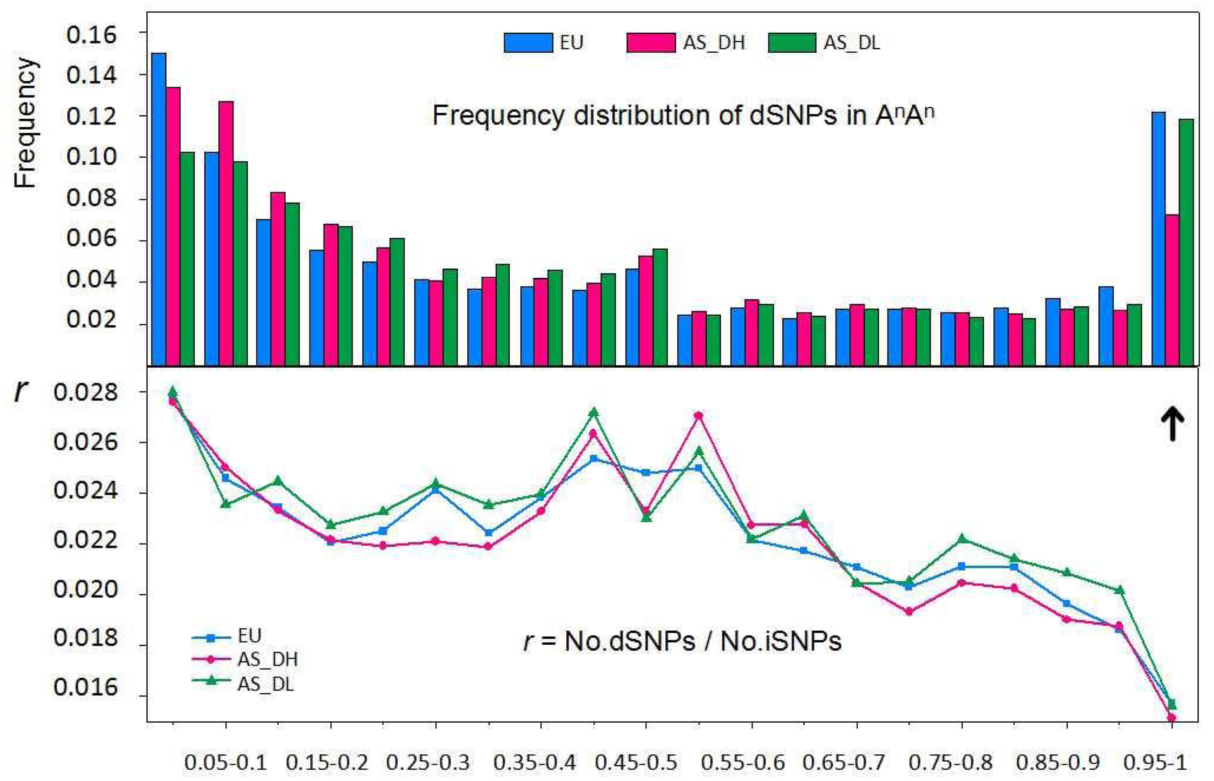
Genetic load estimation for European and Asian groups. Frequency distribution of dSNPs in AA subgenome of European (EU) and two Asian subgroups (AS_DH and AS_DL) (above) with an arrow indicating fixed dSNPs (with 0.95-1 frequency in target population) and relative frequency of deleterious to neutral variants (iSNPs) (below) in the three groups were illustrated.

## Discussion

We have been able to dissect the rapid genome differentiation among different lineages of a complex young domesticated allopolyploid. In addition, we have gained insights to guide further genomic improvement of this recent crop by understanding the pattern of historical genetic improvement. This was facilitated by genome wide comparisons based on *de novo* assembly and population evolution. We generated a high quality *B. napus* genome sequence, rigorously anchored to an improved dense *B. napus* genetic linkage map and with the largest and most complete anchored genome size (892 Mb) to date. This provides an excellent reference for *B. napus*, particularly representing the Asian group, as well as a comprehensive genomic resource for understanding *B. napus* pedigree and breeding. Compared with existing ‘draft genomes’, we have a more complete assembly with improved quality parameters (Figure 1; Table 1, Figure S3-4), which provides an excellent resource for pan-genome dissection in this species. The extensive structural variation between the different genomes representing European, Asian double high and Asian double low crop groups (Figure S6, Table S5-6), enabled us to ask deep questions about the processes underlying rapid genome reorganization in a young allopolyploid species.

We provide genomic evidence supporting a recent origin (<500 years) for Asian rapeseed derived from European germplasm (Figure 3, Figure S11-12), and following introduction from Europe, Asian rapeseed appears to have experienced strong artificial selection, including inter-species (*B. rapa*) and later intra-species introgression events (Figure 2, Figure 4-5; Figure S15). The inter-species introgression caused a significant increase in genomic diversity and decrease of dSNPs accumulated in the Asian group relative to European progenitor material (Figure 6, Figure S14). This was counteracted by increasing dSNP accumulation in the later intra-specific introgressions from European *B. napus* that introduced the key canola quality traits of low seed oil erucic acid and low meal glucosinolate content (Figure 4). Meanwhile, the strong local selection applied to Asian rapeseeds such as appropriate flowering time adapted to the local growing season also increased their LD relative to the European group. Thus, the two historical introgression events separated over ~450 years, combined with local selection re-shaped the Asian genomes. The latter event may be the first report of such a dramatic change in accumulation of deleterious mutations and genomic diversity in polyploid crops resulting from a relatively short breeding process (around 40 years). The results highlight the high plasticity and dynamics of the polyploid rapeseed genome under artificial selection for local adaptation and target traits.

As evident from our analysis of the NY pedigree, breeding programs typically increase genetic load progressively in released cultivars, resulting from selective sweeps that can drag loci linked to deleterious variants to high frequency (Marsden *et al*., 2016). However, for rapeseed a lower dSNP accumulation rate in the Asian group relative to their progenitor (European) group was observed. Our evidence (Figure 6), suggests that the *B. rapa* hybridisation event and subsequent introgression in the Asian group (1950’s) slowed the accumulation of dSNPs in the Asian group. This may be due to the hybrid event (Szadkowski *et al*., 2010) switching the derived deleterious variants of *B. napus* back to an ancestral *B. rapa* state, and greatly reducing the allele frequency of fixed deleterious alleles. We infer that the introgression of *B. rapa* may have made a greater contribution to reducing genetic load in Asian *B. napus* and consequently enhancing relative fitness or local adaptation for Asian rapeseeds. We also observed the significant impact of the *B. rapa* hybrid or subgenome breeding on the genetic background of the allopolypoid *B. napus*. Meanwhile, the analysis suggests new breeding approaches based on purging deleterious mutations from crop genomes.

In conclusion, our study significantly improves our understanding of the young and dynamic domesticated genome of *B. napus*, and in particularly the differentiation and local adaptation of Asian and European rapeseeds. It is apparent that both introgression of the *B. rapa* A genome and subsequent intra-specific crossing between the Asian and European groups has significantly broadened the gene pool of *B. napus* and enlarged the genetic distance of the hybrid parents. In particular, there is scope to target further recombination to the genome regions showing limited and frequent changes.

We conclude that the polyploid origins of *B. napus* confers a very high level of genome plasticity that facilitates rearrangements beneficial for rapid adaption to new environments and local selection. This mechanism, which has evolutionary advantages, is clearly beneficial for crop breeding. In particular, it appears that strategies that re-introduce diploid progenitors contribute to generating novel germplasm in a shorter time span than required for diploid or less complex genomes.

## URLs

BRAD, http://brassicadb.org/brad/;

GENOSCOPE, www.genoscope.cns.fr/brassicanapus/.

## Methods

### Plant materials and phenotypes

Ningyou 7 (NY7), a Chinese semi-winter cultivar with double high seed quality (high glucosinolates and high erucic acid), and its parents in the pedigree were analyzed in this study (Figure 1a). These lines were homozygote. On the purpose of genome sequencing and genetic mapping, NY7 has been initially cultured to a DH line. The other lines were self-pollinated for advanced generations. NY7 was bred from an intraspecific cross between “Ningyou 1” (NY1, elite cultivar in Yangzi River of China in 1960’), and “Chuangyou2” (CY2, elite cultivar in southwest China in 1950’), the latter of which was bred from an interspecific cross between a *B. napus* old popular cultivar “Shengliyoucai” (SL) in China and a Chinese oilseed *B. rapa* cultivar “Chengduaiyoucai” (CDA) as well introduced in previous studies (Bancroft *et al*., 2011; Wang *et al*., 2017b).

In China, there are several accessions with unified name as “Shengliyoucai” (SL) for the imported *B. napus* accessions from Japan or their derived cultivars by pedigree and single plant breeding. To determine the founder parent SL in the NY7 pedigree, four SL accessions were collected (three of the SL accessions provided by the Germplasm Research Group of Chinese Agricultural Academy Oilseeds Research Institute) and all first sequenced to at least 15× coverage (Table S1) and a phylogenetic tree of *B. napus* accessions including the four SL accessions based on their SNPs of AA and CC genomes was constructed (“SL” in Figure S8). The SL accession with the closet to NY7 was renamed as SL-1 as the parental line of NY7 and further deep sequenced for subsequent sequencing and analysis (Table 1).

For elucidating the origin of Asian rapeseeds and population genetic analyses, other *Brassica* accessions with public genomic data by previous studies were included. Finally, a total of 127, 34, and 37 accessions for *B. napus*, *B. rapa*, and *B. oleracea*, respectively, were used in this study (Table S8). The 127 *B. napus* accessions include 59 from European lines, 38 Asian double-high, and 30 Asian double-low rapeseeds.

### Mapping populations and genetic map construction

The genetic mapping population, *Bna*TNDH population by crossing NY7 with an European winter cultivar DH line with double low seed quality, Tapidor (abbreviated as “T”), were used to construct genetic map, which are used as a reference genetic mapping population internationally accumulated with a plenty of genotypes and phenotypes for years (Qiu *et al*., 2006; Zhang *et al*., 2016). Recently, a total of 1,904 consensus quantitative trait loci (QTLs) accounting for 22 traits in up to 19 environments identified in *Bna*TNDH population were summarized using a dense SNP-based genetic map constructed with 182 DH lines (Luo *et al*., 2017). With consideration of multiple trait association with the complex trait, a total of 525 essential QTL were further integrated for dissecting seed yield (Luo *et al*., 2017). The information was referenced in this study. By enlarging the population size, a total of 353 DH lines of the *Bna*TNDH population were further used for genetic mapping genotyped with the Illumina Infinium *Brassica* 60K SNP array in this study, the genotypes and genetic map of 182 DH lines of which has been available previously (Zhang *et al*., 2016). As a result, a total of 2,964 genetic loci including 14,936 SNP markers were mapped in the *Bna*TNDH genetic map (*Bna*TNDH 2.3) (Table S2). All of the markers were filter with high quality parameters with less than 0.05% missing, MAF≥ 0.1. The markers showing the same segregation pattern were classified into the same genetic bins, and one of the markers with the highest quality were choose to construct genetic map using JoinMap 4.0 (Van Ooijen, 2006). Double crosses and recombination frequency were checked by Mapdisto 2.0 (Lorieux, 2012; Zou *et al*., 2014; Zou *et al*., 2016).

### Genome sequencing

The whole NY7 genome sequencing was performed through a combination of sequencing technologies: Illumina sequencing and PacBio Single Molecule Reat Time (SMRT) sequencing. Approximately 200 Gb Illumina sequencing data (170.9× genome coverage) was generated, including three pair end libraries with insertion size of 300, 500, 800 bp and five mate pair libraries with insertion size of 2, 3, 5, 10 and 20 Kb (Table S1). A total of 36 Gb Pacbio sequencing data (30.7× genome coverage) with an average subreads length 7.1 Kb was generated with the PacBio RS II and sequel platform, including 4 SMRT cells from RS-II P5C3 chemistry and 9 SMRT cells from sequel V2 chemistry. Meanwhile, four parental lines in NY7 pedigree (NY1, CY2, SL-1 and CDA) were deep sequenced by the libraries with both short (300 and 800 bp) and large (3 and 10 Kb) insertions, and the additional three SL accessions (SL-2, SL-3, and SL-4) were also re-sequenced to over 15× (Table S1) to represent the early Asian double high accessions.

### HiC experiments and sequencing

The NY7 tissues were fixed in MS buffer (10 mM potassium phosphate, pH 7.0; 50 mM NaCl; 0.1M sucrose) with 1 ml of ~36% formaldehyde solution under vacuum for 30 min at room temperature. After fixation, the leaves were incubated at room temperature under vacuum in MC buffer with 0.15 M glycine for 5 min. Approximately 2 g fixed tissue was homogenized with liquid nitrogen and resuspended in nuclei isolation buffer and filtered with a 40-nm cell strainer. The procedures for enriching nuclei from flow-through and subsequent denaturation were done according to a 3C protocol established for maize (Louwers *et al*., 2009).

The chromatin extraction was similar to that used in the DNase I–digestion experiment. Briefly, chromatin was digested for 16 h with 400 U HindIII restriction enzyme (Takara) at 37 °C. DNA ends were labeled with biotin and incubated at 37 °C for 45 min, and the enzyme was inactivated with 20% SDS solution. DNA ligation was performed by the addition of T4 DNA ligase (Fermentas) and incubation at 16 °C for 4~6h. After ligation, proteinase K was added to reverse cross-linking during incubation at 65 °C overnight. DNA fragments were purified and dissolved in 86 µL of water. Unligated ends were then removed. Purified DNA was fragmented to a size of 300–500 bp, and DNA ends were then repaired. DNA fragments labeled by biotin were finally separated on Dynabeads^®^ M-280 Streptavidin (Life Technologies). The HiC experiments procedures were similar to those described previously in arabidopsis and cotton (Wang *et al*., 2017a). HiC Libraries were controlled for quality and sequenced on an Illumina Hiseq X Ten sequencer.

### *De novo* assembly for scaffolds

To avoid systematic errors from sequencing reads, the raw Illumina reads were filtered using NGSQC v2.3.3 (Patel and Jain, 2012) and corrected using Lighter (Song *et al*., 2014) with default setting. Firstly, these qualities filtered and corrected Illumina data were assembled using SOAPdenovo2 (Luo *et al*., 2012). Secondly, the clean reads were reused to improve the SOAPdenovo2 assembly by filling the gaps using Gapcloser v2.1 (Luo *et al*., 2012) and extending the scaffolds using SSPACE v1.0 (Boetzer *et al*., 2011) and OPERA-LG v2.1 (Gao *et al*., 2016). Thirdly, the Pacbio reads corrected by clean Illumina reads with software LoRDEC 0.6 (Salmela and Rivals, 2014) were used to fill the gaps by PBjelly (English *et al*., 2012). As a result, the NY7 assembly was represented by 9,898 scaffolds including 993.9 Mb with high assembly quality of a scaffold N50 1.97 Mb and a contig N50 44.6 Kb.

### Correction of chimeric scaffolds

According to the dependable synteny relationship of NY7 assembly with three linkage maps and A^r^ and C° subgenomes, 180/9,898 chimeric scaffolds that mapped to multiple linkage groups were identified. In these chimeric scaffolds, the breakpoint regions were confirmed with at least three matches to different linkage groups and then the breakpoints were set to the largest gap if there existed gaps in the breakpoint regions or were set in the middle of breakpoint regions. As a result, a total of 180 chimeric scaffolds were corrected and NY7 assembly could be improved with a scaffold N50 1.27 Mb and a contig N50 44.0 Kb.

### Super-scaffolding with HiC data

Three-dimensional (3D) chromosome conformation capture sequencing (HiC) was further performed to improve the assembly. Approximately a total of 100 coverage Illunima pair-end reads of NY7 genome was yielded. We mapped the HiC data to the assembly using Bowtie v2.2.1 (Langmead and Salzberg, 2012) with extra parameters ‘--reorder’ and ‘--very-sensitive’ to force SAM output to match order of input reads and yield high precise result, respectively. The software Samtools (Li *et al*., 2009) was used to manipulate the BAM file and remove potential PCR duplicates. Then we used Lachesis (Burton *et al*., 2013) to cluster, order and orientate the scaffolds and created the raw HiC assembly with the mapping result in the last step, but the raw Lachesis HiC assembly would produce super large scaffolds even larger than the actual chromosome size. For the assembly accuracy, the HiC mapping links matrix was generated and custom scripts were used to find the weak point among the HiC assembly and split the raw HiC assembly. As following, the corrected HiC assembly was aligned to the three linkage maps and A^r^ and C° subgenomes to remove the abnormal synteny relationship. Finally, we got the NY7 HiC assembly with a scaffold N50 6.91 Mb and a contig N50 44.0 Kb.

### Pseudomolecule construction

The backbone linkage map, *Bna*TNDH 2.3 (Table S2), and two other published linkage maps (DYDBAA (Chalhoub *et al*., 2014) and BS (Delourme *et al*., 2013)), were used to construct the pseudomolecules and a final set of 12,825 unique markers were utilized to anchored the HiC scaffolds using blast+ 2.3.0 with parameters “-evalue 1e-10”, only the markers with best hits having no same score in the NY7 genome can be defined as unique markers. Then we constructed 19 pseudo-chromosomes using Allmaps (Tang *et al*., 2015), covering 890 Mb. As a result, 89.5% of assembled genome size and 828 Mb (93%) were oriented.

### Improving the assembly of NY7 with gene models constructed with the transcriptome of *Bna*TNDH population

Illumina mRNAseq reads from 45 lines of the Tapidor and NY7 double haplotype (*Bna*TNDH) mapping population and 2 parental lines (NY7 and Tapidor) were mapped with this genome reference using methodology developed and deployed previously (Bancroft *et al*., 2011). The SNP scoring strings were filtered to retain only simple SNPs (i.e. polymorphisms between resolved bases). To avoid minor allele being unresolved base, a SNP is flagged as simple when the most abundant two alleles scored are simple. After filtering homozygous SNPs between parental lines Ningyou7 and Tapidor, the remaining SNPs are displayed in genome sequence order as genome-ordered graphical genotypes (GOGGs). Alleles that are inherited from NY7 are coloured in light red and those inherited from Tapidor in dark blue (Table S3).

We developed a new NY7 genome sequence resource by cutting and inserting 54 segments from the draft genome sequence, based on the genome-ordered graphical genotypes (GOGGs) (Table S3). The re-assembly of the split segments was designed based on the congruence of genotypes of the mapping population. The new NY7 genome sequence was re-assembled based on an automation of concatenating of the sequence segments. Another iteration of SNP scoring and GOGGs generation from this resource showed the improved congruence of genotypes (Table S3).

### Assembly confirmation with HiC contact maps

~100X HiC coverage illumina were remapped to the 19 pseudo-chromosomes and normalized using HiC-Pro 2.10.0 (Servant *et al*., 2015) with parameters “FILTER_LOW_COUNT_PERC=0 and BIN=100000” and the HiC contact maps with a bin 100 Kb generated by HiC-Pro was used to plot HiC contact heatmaps for whole 19 pseudo-chromosomes using HiCPlotter 0.7.3 (Akdemir and Chin, 2015) with parameters “-tri 1 -wg 1 -o WholeGenome” (Figure S3).

### QTL mapping in NY7 and Tapidor assembly

Consensus QTL for different traits and essential QTL for seed yield from Luo *et al*. (2017) were mapped to NY7 assembly using BLAST, and unique markers were used to determine the location of these QTLs. After removing abnormal markers of other pseudo-chromosomes or distant location from most markers, 1526 consensus QTLs and 303 essential QTLs larger than 1 Kb were identified.

As Tapidor assembly was only in scaffold level, reads mapping and synteny comparison between two assemblies were applied to detect the variance of QTLs in NY7 and Tapidor assembly. Bowtie2 with default settings were used to align Tapidor reads to NY7 assembly and sambamba v0.6.5 with ‘depth’ module was used to calculate the depth of 303 essential QTL regions in NY7 assembly. Nucmer program form Mummer 3.23 package was used to detect the co-linearity between NY7 and Tapidor assembly, only synteny blocks with identity >-90% will be considered in the comparison and the accumulated length of synteny blocks in the QTL region will be applied in the next stage. Ribbon (Nattestad *et al*., 2016) and JBrowse 1.13.1 (Buels *et al*., 2016) were applied to visualize the compassion results. Finally, the overlaps between top 15% QTL loci with most minimum reads coverage and top 15% QTL loci with most minimum assembly similarity identity were defined as the distinct sequence between NY7 and Tapidor assembly.

### Genome gene and repeat annotation

We built the *de novo* repeat library from the assembled genome using RepeatModeler (Chen, 2009). A total of 352.8 Mb repetitive elements covering 41.29% of NY7 genome were identified using RepeatMasker 4.0.8 (Chen, 2009) with default settings. *De novo* gene structure predictions were carried out using AUGUSTUS 3.2.2 (Stanke *et al*., 2006), GeneMark.hmm (Lukashin and Borodovsky, 1998), and FGENESH 2.6 (Salamov and Solovyev, 2000). The coding sequences of *A. thaliana* (http://plants.ensembl.org/Arabidopsis_thaliana/, TAIR10),*B.rapa* (http://plants.ensembl.org/Brassica_rapa/,IVFCAASv1), *B. napus* (Darmor-*bzh* (http://www.genoscope.cns.fr/brassicanapus/data/,v5)andZS11 (http://ocri-genomics.org/Brassia_napus_genome_ZS11/,V201608) and *B. oleracea* (http://plants.ensembl.org/Brassica_oleracea/,v2.1) were downloaded to perform the homology evidence predictions using gmap (Wu and Watanabe, 2005). A total of ~20Gb RNA-seq data were obtained from Bancroft *et al*. (2011), which included the data from four *Bna*TNDH lines, T, NY7 and the parents in the pedigree of T and NY7. We aligned the RNA-seq reads to the NY7 assembly using Tophat 2.1.1 (Trapnell *et al*., 2009). A set of assembled transcripts were obtained using Cufflinks 2.2.1 (Trapnell *et al*., 2010) with default settings. Meanwhile, the RNA-seq data were assembled using Trinity 2.4.0 (Grabherr *et al*., 2011) with default settings. The assembled transcripts from Cufflinks and Trinity were integrated using PASA 2.0.2 (Haas *et al*., 2003) pipeline to the RNA-seq evidence for gene predictions. All gene models from the evidence above were combined using EVM (Haas *et al*., 2008) with higher weight in the evidence of RNA-seq results. The final gene set yielded 104,179 genes, and the majority (97.0%) had EST mapping evidence support.

### Detection of small variations (SNP and Indel)

The raw paired-end reads were first filtered into clean data using NGSQCtookit v2.3.3 (Patel and Jain, 2012). The cut-off value for PHRED quality score was set to 20 and the percentage of read length that met the given quality was 70. Clean reads of each accession were mapped to our assembled NY7 reference genome and Darmor-*bzh* genome as well using Bowtie v2.2.1 (Langmead and Salzberg, 2012) with default settings. Consecutive steps using Samtools v0.1.19 (Li *et al*., 2009) and GATK v2.3 (McKenna *et al*., 2010) were applied for variants detection. Potential PCR duplicates were removed by ‘Samtools rmdup’. Alignments around small indels were remapped with ‘IndelRealigner’, and raw variants were called based on the realigned bam file. Using the called variants as known sites, ‘BaseRecalibrator’ and ‘PrintReads’ in the GATK were applied for base-pair scores recalibration. The proceeded BAM files of each sample were used for the multi-sample variant genotyping. ‘UnifiedGenotyper’ in GATK was applied to generate the raw variant calls with parameters “-stand_call_conf 30”. To reduce the variants discovery rate, the SNP calls were filtered according to the following threshold: QUAL < 30, DP < 10, QD < 2. Potential variant annotation and effect were predicted by SnpEff v3.6 (Cingolani *et al*., 2012).

### Detection of structure variations

The structure variations between NY7, Darmor-*bzh* and other genomes were detected by svmu (Chakraborty *et al*., 2018), after each NY7 chromosome was aligned to the corresponding chromosome of Darmor-*bzh* by mummer (nucmer –mumreference --noextend). To detect specific genomic regions in NY7 in contrast to Darmor-*bzh*, we first mapped reads of NY7 to Darmor-*bzh* genome, and extract orphan reads that could not be mapped to Darmor-*bzh* genome. The orphan reads were relocated to NY7 genome. Genomic regions (≥ 100 bp) that were mapped by at least two orphan reads per base were considered NY7 specific genomic regions. The specific regions of Darmor-*bzh* compared to NY7 were identified using the same approach.

### Detection of identity by decent (IBD) blocks in the pedigree

For determining the IBD origin of NY7 genomic segment, we took advantage of our assembled genomes of NY7 pedigrees. NY7 genome data was first divided into blocks with a length of 50 Kb. All pedigree scaffolds were taken as a reference genome database for tracing sequence origin of NY7. MUMmer3.23 (Kurtz *et al*., 2004) was applied to do the 1-to-1 alignment. We calculated pedigree’s match length in each 50 Kb block, and assigned the origin of the block if a pedigree has longest accumulated alignment length. The demonstration of IBD was draw with Perl SVG module in each chromosome.

### Phylogenetic and principal component analysis

We filtered SNPs with VCFtools with parameters “-maf 0.01 -max-missing 0.9” for *B. napus* accessions used in this study (a total of 127 lines, see Table S8), phylogenetic tree was constructed using Fasttree (Price *et al*., 2009) with 1,000 replicates for bootstrap confidence analysis. MEGA v7 (Kumar *et al*., 2016) was applied to draw the constructed tree. Principal component analysis was performed by SNPRelate v0.9.19 (Zheng *et al*., 2012).

### Demographic analysis

To minimize bias in demographic analyses due to selection, SNPs in intergenic regions were used. The best parameters for fitting model were estimated by ∂a∂i v1.6.3 (Gutenkunst *et al*., 2009). The alleles were down-sampled via hypergeometric projection for each group. Folded spectrum was used for each pool. Four demographic models were considered for each type, i.e. (1) EU and AS independently split from progenitors (*B. rapa* or *B. oleracea*) Br/Bo, and EU earlier than AS; (2) EU and AS independently split from Br/Bo, and AS earlier than EU; (3) EU split from Br/Bo, and AS split from EU; (4) AS split from Br/Bo, and then EU split from AS (Figure S10). For inference of Asian double-low rapeseeds, five demographic models were considered, i.e. (1) ASL and ASH independently split form EU, and AS_DH earlier than AS_DL; (2) AS_DL and AS_DH independently split form EU, and AS_DL earlier than AS_DH; (3) AS_DH split from EU, and then AS_DL split from AS_DH; (4) AS_DL split from EU, and AS_DH split from AS_DL; (5) AS_DH split from EU, and then AS_DL split from AS_DH, EU has introgressed to AS_DL (Figure S11). Different demographic models were compared with the basis of the relative log likelihoods of the models given the observed site frequency spectrum. Thirty independent runs with randomized starting points were executed for each candidate model, and the average value was chosen based on the best fitting parameters.

Effective population size and split timing between EU and AS populations was also inferred with SMC++, which analyzes multiple genotypes without phasing (Terhorst *et al*., 2017). The evolutionary rates (9e-9) were used for age estimates (Qi *et al*., 2017).

### Population parameters estimation

The genome was scanned in a 100 Kb window size and the population parameters (*π*, *F*_ST_) were estimated for each window by VCFtools (Danecek *et al*., 2011). Nucleotide diversity (*π*) was measured with parameters ‘--window-pi 100,000 --window-pi-step 10,000’. The average 100k window size π value was taken as the genetic diversity. For measurement of population differentiation, *F*_ST_ was calculated with the setting ‘--fst-window-size 100000 --fst-window-step 10000’. *Z*-transformation was applied to locate divergent regions from the extreme tails by applying a threshold of 3 standard deviations. The non-reductant genes residing in these regions were taken as putatively divergent genes between different populations.

### Local ancestry inference

Local ancestry inference was performed by Loter (Dias-Alves *et al*., 2017), which can use just haplotype data to infer ancestry across the chromosomes of an admixed individual from two proposed ancestor populations. In this study, the Asian double-low rapeseed population was employed as the descendant population, and European rapeseed and Asian double-high rapeseed population were employed as the two parental populations. For inference of potential *B. rapa* introgression in Asian rapeseed, twelve Asian rapeseeds for oil use and European rapeseed were employed as parental populations.

### Identification of deleterious mutations

Deleterious SNPs (dSNP) were predicted by SIFT (Kumar *et al*., 2009). The combined *B. rapa* and *B. oleracea* genomes (EnsemblePlants, release-37) were used for building reference database, considering using either Asian (NY7) or European (Darmor-*bzh*) rapeseed genome as reference would substantially cause reference bias for dSNP identification. A dSNP was defined if the value calculated by SIFT for a SNP had a normalized probability < 0.05. To calculate the site frequency spectra for Asian and European populations, the number of deleterious alleles in a population was calculated as twice the number of homozygous variants plus the number of heterozygous variants.

## Acknowledgements

This work was supported by the National Basic Research Program of China (2015CB150200), the National Key Research and Development Program for Crop Breeding in China (2016YFD0100305, 2016YFD0101300), the 111 Project (B17039) and Jiangsu Collaborative Innovation Center for Modern Crop Production. This work was supported by UK Biotechnology and Biological Sciences Research Council (BB/L002124/1), including work carried out within the ERA-CAPS Research Program (BB/L027844/1). We acknowledged Prof. Xiaoming Wu of Chinese Agricultural Oilseed Research Institute for providing the three Shengliyoucai accessions.

## Author contributions

L.F. and J.M. conceived and designed the project. J.Z. and W.M. collected materials, performed the experiments, developed genetic maps, QTL analysis, and result analysis. L.M. and J.Q. performed genome assembly. J.Q., L.M., L.J., D.W., N.S., Y.S., J.Z., M.W., R.L., D. H., M.C., C.Y., and L.F. managed sequencing and analyzed the data. Z.H. and I.B did GOCGs for genome assembly corrections. J.M., I.B., L.S., and C.Y. discussed the data. J.Z., J.Q., L.M., and L.F. wrote the manuscript. G.K., I.B., and J.M. revised the manuscript.

## Supporting information

Additional Supporting Information may be found online in the supporting information tab for this article:

Table S1 Summary of the sequenced genomic data in this study.

Table S2 The linkage genetic map of *Bna*TNDH mapping population version *Bna*TNDH 2.3.

Table S3 Re-assembling of the NY7 genome according to the genome-ordered graphical genotypes constructed using *Bna*TNDH population.

Table S4 Genome completeness assessment using CEGMA and BUSCO.

Table S5 Number of different types of genomic variations for NY7 compared with other *de novo* assembly genomes.

Table S6 Specific genomic regions above 5 Kb between NY7 with other *de novo* assembly genomes.

Table S7 QTL regions with distinct sequence diversity between Tapidor and Ningyou7 assembly.

Table S8 *Brassica* accessions used in this study.

Table S9 Details of demographic model for the origin of Asian rapeseeds.

Table S10 Divergent regions and QTLs located between the Asian and European cultivars.

Table S11 The conserved regions introgressed from *B. rapa* in Asian double-high rapeseeds and QTL located.

Table S12 QTLs in IBD genomic blocks of NY7 pedigrees.

Table S13 Nonsynonymous SNPs in flowering time genes with high allele frequency differentiation between Asian and European populations potentially due to *B. rapa* introgression.

Figure S1 The pedigree of NY7.

Figure S2 Genomic survey of NY7 and its pedigrees with *K*-mer distribution.

Figure S3 The HiC contact maps for NY7 whole genome and NY7 chrA03.

Figure S4 The collinearity between three *B. napus* genetic maps and the “NY7” genome assembly.

Figure S5 The pipeline for genome assembly and pseudo-chromosome construction for NY7.

Figure S6 The genomic synteny between NY7 and other *de novo* assembly of *B. napus* and two diploid progenitors. a-e present the synteny between NY7 and Darmor-*bzh*, progenitor diploid *B. rapa* and *B. oleracea*, Tapidor published in Bayer *et al*. 2017, Tapidor updated in this study, and ZS11, respectively, while f presents the picture of the synteny between these two Tapidor assembly versions.

Figure S7 Demonstration of the QTL located in the genomic divergent region between the NY7 and Tapidor genome. (A) Genomic alignment view of Tapidor assembly to NY7 reference on chromosome C03. All of the QTL identified in C03 were labeled with ‘es.’ at the top the rectangle. The arrow shows a QTL (*es. C3-3*) region with present and absent sequence variation between Tapidor and NY7. (B) Significant different reads coverage to the NY7 genome within the QTL (*es. C3-3*) region between Tapidor (low box) and NY7 (upper box).

Figure S8 Phylogenetic trees for *B. napus* accessions based on SNPs of AC genomes. The EU, AS_DL, AS_DH present the European rapeseed (*B. napus*), Asian rapeseed (*B. napus*) with double low (low erucic acid and low glucosinolates) and double high seed quality traits, respectively.

Figure S9 Genetics diversity (*π*) of different chromosomes in different populations. The EU, and AS present the European rapeseed (*B. napus*), and Asian rapeseed (*B. napus*), respectively. The EU, AS_DL, AS_DH present the European rapeseed, Asian rapeseed with double low and double high seed quality traits, respectively.

Figure S10 Linkage disequilibrium of European and Asian rapeseed populations.

Figure S11 Demographic models for the origin of Asian rapeseeds.

Figure S12 Demographic models for the origin of Asian double-low rapeseeds.

Figure S13 A genomic view for the reads mapping information for *CDF2* homolog in the lineage of NY7.

Figure S14 The frequency distribution of deleterious SNPs (dSNPs) and neutral intergenic SNPs (iSNPs) in (a) the NY7 pedigree (b) CnCn genome of Asian and European populations.

Figure S15 A summary for the origin and breeding cycle of Asian rapeseeds and the dynamic changes of genomic diversity and deleterious SNP (dSNP) accumulations during the Asian local breeding programs. The Asian genomes were significantly re-shaped by two genetic introgression events by *B. rapa* and double low European *B. napus*.

